# Proximate and Genetic Analysis of Blackfin Tuna (*T. atlanticus*)

**DOI:** 10.1101/2020.11.03.366153

**Authors:** Yuridia M. Núñez-Mata, Jesse R. Ríos Rodríguez, Adriana L. Perales-Torres, Xochitl F. De la Rosa-Reyna, Jesús A. Vázquez-Rodríguez, Nadia A. Fernández-Santos, Humberto Martínez Montoya

## Abstract

The tuna meat is a nutritious food that possesses high content of protein, its low content of saturated fatty acids makes it a high demand food in the world. The *Thunnus* genus is composed of eight species, albacore (*T. alalunga*), bigeye (*T. obesus*), long tail tuna (*T. tonggol*), yellowfin tuna (*T. albacares*), pacific bluefin tuna (*T. orientalis*), bluefin tuna (*T. maccoyii*), Atlantic bluefin tuna (*T. thynnus*) and blackfin tuna (*T. atlanticus*). The blackfin tuna (BFT) (*Thunnus atlanticus*) represent the smallest species within the *Thunnus* genus. This species inhabits the warm waters of the West Atlantic Ocean, from the shore of Massachusetts in the north, to Rio De Janeiro in Brazil. The first objective of this study was to evaluate the nutritional composition of BFT captured in the Gulf of Mexico, we determined ash, moisture, fat, protein and carbohydrates in BFT muscle and compared the obtained data with the nutritional reports from commercial tuna species including yellowfin tuna, Atlantic bluefin tuna and salmon (*Salmo salar*).

Secondly, we report the genetic diversity and genetic differentiation of BFT within its geographical distribution range using the Cytochrome Oxidase I (COI) and control region sequenced data and from specimens collected in the Gulf of Mexico. We observed a nucleotide diversity π=0.001, 24 segregating sites and 10 parsimony informative. Within the CR we found nine different haplotypes π=0.044, 39 segregating sites, 16 parsimony informative sites. We concluded that according with the haplotype distribution there are differences among the BFT from the Gulf of Mexico and the North Atlantic compared to the South Atlantic. The Caribbean Sea is a migration point of the BFT, where all except the South Atlantic haplotypes were found.

## 1. INTRODUCTION

According to the FAO, the species within *Thunnus* genus are economically relevant and in general are overexploited. However, currently, only six out of eight species of the genus *Thunnus* are commercially exploited around the world. According to their phenotypic characteristics, the *Thunnus* genus is subdivided into two subgenera. The first group, the *Thunnus* subgenus (blue tunas) is composed of the species albacore (*T. alalunga*), Atlantic blue (*T. thynnus*), Pacific blue (*T. orientalis*) and south blue tuna (*T. maccoyii*). Secondly, the *Neothunnus* subgenus (tropical yellow tunas) is composed of the blackfin tuna (*T. atlanticus*), yellowfin tuna (*T. albacares*), longtail tuna (*T. tonggol*) and bigeye tuna (*T. obesus*) (Collete and Nauen, 1983, Díaz-Arce et al., 2016, Gong et al., 2017, Viñas and Tudela, 2009)

Commercial exploitation of blackfin tuna (BFT) is not yet wholly developed (Doray et al., 2004), However, this is the only tuna species used for either artesian fisheries or sport fishing. Its body size is in average 100 cm, with an average body weight between 5 to 7 kg and a biological cycle of 5 years. The species inhabits the warm waters of the Western Atlantic, from Massachusetts to the shore of Rio de Janeiro in Brazil, including the Gulf of Mexico (GdM) and the Caribbean Sea.

In the Gulf of Mexico, the BFT stands out in local trading. Several studies of traceability in Mexico recognized that the meat products of BFT are commercialized in local markets as generic tuna. Therefore, there are not detailed descriptions about the amount of BFT commercialized in Mexico. A study of incidental catching of blackfin tuna as a secondary species during the yellow fin tuna fishing reports 137 tons in a ten year-period (Ramirez-López, 2016). Unlike other tuna species, the genetic diversity, genetic stocks and nutritional properties of BFT remains poorly known. Genetic structure studies in BFT found differences between the population from GOM and the Northwest Atlantic as well migration patterns from the GOM to the Atlantic, a spawning behavior and ocean currents in the GOM may explain these findings (Saxton, 2009). However, to date very little is known about this species at the level of the population and subpopulation structure throughout its distribution range. The traceability of the eight species of tunas is commonly conducted with the Cytochrome B (Cyt B), Cytochrome Oxidase I (COI) or the control region (CR), which are the most widely accepted markers for the species identification (Alvarado Bremer et al., 1997, Alvarado Bremer et al., 2005, Chow and Kishino, 1995, Chow et al., 2000, Durand et al., 2005, Hebert et al., 2003, Kochzius et al., 2010, Niwa et al., 2003, Sarmiento-Camacho et al., 2018, Viñas and Tudela, 2009, Ward et al., 2005). Similar results with mitogenomes and genotyping by sequencing techniques review the utility of additional markers and revisited the COI gene as barcoding for tuna species. Particularly, the COI gene of BFT is represented in the BOLD system platform in different stages of its life cycle (larva and adult) and in almost its complete range of distribution, since Florida in the US to Sao Paulo in Brazil. The aim of this study was to evaluate the nutritional composition of BFT captured in the Gulf of Mexico and identify the genetic diversity of the species within its range of distribution.

## 2. MATERIALS AND METHODS

### 2.1 GdM BFT proximate analyses

#### 2.1.1 Biological samples

Five BFT specimens were collected in the shore of Matamoros municipality, state of Tamaulipas, Mexico by local fishermen in July, 2018. The fish were transported in refrigeration, once in the lab, the samples were measured, weighted and stored at −20 °C for further analyses. BFT collected in the GOM were subjected to in vitro digestibility and proximate analysis to evaluate its nutritional content following the procedures described by the Association of Official Analytical Chemists (AOAC, 2012). In this study, we determined moisture, ash, fatty acids, protein and carbohydrates from homogenized muscle.

#### 2.1.2 BFT proximate analysis

The proximate analyses were carried out based on official Association of Official Analytical Chemists methods (AOAC). The total protein content was determined using Dumas’s method (official method 968.06), along with moisture (official method 14.003), ash (official method 14.006), crude fat (official method 7.056) and crude fiber (official method 962.09). Carbohydrates were calculated by difference.

#### 2.1.3 BFT in vitro digestibility

The human in vitro digestion analysis was performed according to the procedure described by Hsu et al. (1977) and modified by (Soon-Mi et al., 2010, Xing et al., 2017). Five grams of dry BFT muscle were weighted and dissolved in 100 mL of distilled water heated to 37 °C. The initial gastric phase was recreated adding 432 mg of porcine pepsin (Sigma Aldrich, Cat. No. P7000-100G) dissolved in 2.7 mL of HCl (0.1N). Three10 mL aliquots were taken, the first after 5 minutes of pepsin addition (GP0), after 30 minutes (GP30) and after 60 minutes (GP60).

The intestinal phase begun after GP60 adding 40 mg of porcine pancreatin (Sigma Aldrich, Cat. No. P1750-100G) and 250 mg of porcine bile (Sigma Aldrich, Cat. No. B8631-100G) previously dissolved in 10 ml of NaHCO_3_ (0.1N). We sampled aliquots at 0 (IP0), 30 (IP30), and 120 (IP120) minutes respectively. All samples were kept at −20 °C until spectroscopy analyses.

The percentage of digested protein was determined by the Biuret method measuring absorbance by duplicate at 540 nm. % of digested protein was calculated by:

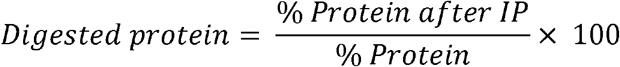

#### 2.1.4 Chromatography analysis of fatty acids

Purification of fatty acids was performed using the fatty acids purification kit cat MAK174 (Sigma Aldrich, St Louis, MO) from 0,15 grams of BFT flour. Aliquots of 100 uL of the total lipid extract obtained were transferred to 2 mL vials and dried under nitrogen flow. 1 mL of boron trifluoride-methanol and 0.3 mL of HPLC grade hexane was added to each vial. The mixture was transferred to 13×100 mm test tubes and heated one hour at 95 °C. Once the tubes were at RT, 1 mL of hexane and 1 mL of distilled water were added to the tubes and vortexed. Tubes were centrifuged (500 g/ 5 minutes) and the supernatant was transferred to a 2 mL vial. For fatty acid analysis, 1 uL of the aliquots was injected to the gas chromatograph (Aligent Technologies, Santa Clara, CA) equipped with an HP88 capillary column (100m x 25mm x 0.2 um) and a flame ionization detector.to perform the chromatographic separations. The carrier gas was helium and the injection Split 100:1. The identification of the FAMEs was carried out by comparison in the retention times of the peaks in the sample with the Supelco 37 FAME mix pure standard by Sigma Aldrich (product number CRM47885), from which a calibration curve was created to calculate the concentration of fatty acids.

### 2.2 BFT population analyses

#### 2.2.1 Data mining

DNA sequences used in this study were mined from BOLD system (Barcode of Life Data System) and from the NCBI (National Center Biotechnology Information) identifying country or region and geographical coordinates of each sample.

For COI and CR we conducted a multiple alignment in the platform MAFFT online (Katoh et al., 2009, Katoh and Standley, 2013) and sequences were manually edited using UNIGENE (Okonechnikov et al., 2012).

#### 2.2.2 COI and CR amplification

The first dorsal fin of each BFT was dissected and DNA was purified from 25 mg of fresh tissue, following the procedure described in the commercial kit DNeasy Blood and Tissue (Qiagen, Valencia CA). The DNA was amplified by PCR and sequenced using the specific primers COI, FishF1 (5’-TCAACCAACCACAAAGACATTGGCAC-3’) and FishR1 (5’-TAGACTTCTGGGTGGCCAAAGAATCA-3’) (Viñas and Tudela, 2009, Ward et al., 2005) and the CR primers Tatlan-F (5’-TACCCCTAACTCCCAAAGCTA-3’) and Tatlan-R (5’-CGAGATTTTCCTGTTTACGGG-3’) designed in this study. The PCR were carried out in a total volume of 12.5 uL using approximately 30 ng of DNA as a template. Additionally, PCR reactions contained 1.24 units of *Taq* polymerase, 1 uM of each primer and 1X of reaction buffer (Promega, Madison WI). Thermal cycles were set up to an initial denaturing step of 95 C, followed by 34 cycles of denaturing at 95 C for 30s, annealing of 52 C for 45s and extension of 72 C for 1 minute. PCR products were purified and sequenced using an ABI PRISM 3130 Genetic Analyzer (Applied Biosystems, Foster City, CA) at the Centro de Biotecnología Genómica Sequencing Core (Reynosa, Mexico) following the manufacturer recommendations.

#### 2.2.3 Genetic diversity analysis and haplotype network

We conducted a nucleotide diversity analysis (π) for COI and CR, the number of segregating sites (S), number of shared and unique haplotypes and the number of parsimony sites among the analyzed sequences in PopArt (Leigh and Bryant, 2015). To analyze demographic aspects of BFT, we calculated the neutrality estimator Tajima’s D.

#### 2.2.4 Analysis of Molecular Variance (AMOVA)

For COI, we performed an AMOVA in Arlequin (Excoffier and Lischer, 2010) testing a three level structure. In the first analysis, we compared all the five geographical zones. In the second we compared only Atlantic-South against the remaining five zones concatenated in one single group. Lastly, in the third structure comparing the found haplotypes. For the CR, we tested one structure comparing the haplotypes found in the Gulf of Mexico with the sequences obtained from NCBI.

#### 2.2.5 Neothunnus phylogenetic analysis

We estimated the phylogenetic relationships among the *Neothunnus* subgenus using the GoM sequences plus the remaining sequences obtained from data mining for COI (n=147) and the CR (n=111). Additionally, we included 18 sequences of *T. albacares*, 4 *T. obesus* sequences and 12 *T. tonggol*. As an external group, we used one sequence from *T. maccoyii*.

The phylogenetic trees were reconstructed using MrBayes in CIPRES (Miller et al. 2011), COI and CR trees were summarized in SumTrees (Sukumaran and Holder, 2008). Consensus trees were visualized and edited using FigTree v1.4.3 (Rambaut, 2012).

## 3. RESULTS

### 3.1 GdM BFT proximate analyses

#### 3.1.1 BFT samples

Every individual sample was identified following biological keys and species was confirmed through the amplification and sequencing of the COI gene. We measured the five specimens captured in the Gulf of Mexico, on average the individuals were 74 ±5.8 cm of total length and 6.25±0.9 kg of weight (Table 1).

**Table 1.** Morphological measurements of analyzed BFT.

| Sample | Length | Total length | Pectoral fin length | Caudal fin length | Wide length | Weight |
| --- | --- | --- | --- | --- | --- | --- |
| BFT1 | 70.0 | 74 | 17.0 | 21.2 | 32 | 6.3 |
| BFT2 | 64.0 | 68 | 17.5 | 22.0 | 20 | 5.5 |
| BFT3 | 73.0 | 83 | 17.5 | 25.0 | 24 | 7.4 |
| BFT4 | 67.5 | 70 | 17.0 | 22.0 | 18 | 5.1 |
| BFT5 | 72.0 | 75 | 17.9 | 23.0 | 22 | 6.9 |
| Mean±SD | 69.3±3.6 | 74±5.8 | 17.4±0.3 | 22.6±5.4 | 23.2±5.4 | 6.3±0.9 |

#### 3.1.2 Proximate analyses

We determined moisture, ash, protein, fat and crude fiber on BFT dry samples in both dry and wet basis conditions (Table 2, Table S1). Moisture content was in a range of 71.65-75.95% and remained constant among the analyzed samples. Ash content on a dry basis was estimated between 4.54 and 6.04%, whereas in a wet basis the range was 1.15-1.55%, although the ash content was similar among the analyzed samples, BFT2 exhibited the highest content of ash (p<0.05) in both dry and wet basis. We observed that BFT have a high protein content on both a dry basis (above 80%) and wet basis (22-24.59%). Remarkably, the biggest specimen, BFT3 showed the highest content in fatty acids on a dry basis (2.24±0.45) and wet basis (0.63 ±0.12) compared with the other analyzed samples.

**Table 2.** BFT dry basis proximate analyses (%).

| Sample | Moisture | Ash | Protein | Fat | Crude fiber | Carbohydrates |
| --- | --- | --- | --- | --- | --- | --- |
| BFT1 | 75.95±0.40<sup>a</sup> | 4.82±0.11<sup>a</sup> | 91.51±1.09<sup>a</sup> | 1.42±0.17<sup>a</sup> | ND | 2.25 |
| BFT2 | 74.11±0.47<sup>bd</sup> | 6.04±0.79<sup>b</sup> | 91.84±4.95<sup>a</sup> | 1.18±0.11<sup>a</sup> | ND | 0.94 |
| BFT3 | 71.65±0.35<sup>c</sup> | 4.54±0.20<sup>a</sup> | 86.78±0.48<sup>a</sup> | 2.24±0.45<sup>b</sup> | ND | 6.44 |
| BFT4 | 75.18±0.11<sup>ab</sup> | 4.56±0.14<sup>a</sup> | 90.32±0.42<sup>a</sup> | 0.88±0.22<sup>a</sup> | ND | 4.24 |
| BFT5 | 75.50±0.57<sup>d</sup> | 4.63±0.22<sup>a</sup> | 92.69±0.56<sup>a</sup> | 0.13±0.07<sup>a</sup> | ND | 2.55 |

Columns with different letters present a significant difference, according to Tuckey's test ( $p < 0.05$ ). ND: no detected by the method.

#### 3.1.3 Determination of fatty acids in BFT

Total content of saturated fatty acids (SFA) was 18.59±4.14, monounsaturated fatty acids (MFA), 35.82 ± 6.94 and polyunsaturated fatty acids (PUFA) 30.93±7.07. The most abundant SFA found in BFT is the stearic acid. Among the unsaturated fatty acids, the MFA palmitoleic acid was found the most abundant, whereas within the PUFA, the decosahexaenoic acid (DHA) was found in the highest amount (Table 3).

**Table 3.** Percentage of fatty acids found in BFT detected by chromatography.

|  |  | BFT1 | BFT2 | BFT3 | BFT4 | BFT5 | Average |
| --- | --- | --- | --- | --- | --- | --- | --- |
| SFA | Total | 23.38 | 21.66 | 17.18 | 17.96 | 12.78 | 18.59±4.14 |
|  | C14:0 | 3.61<sup>a</sup> | ND | 0.17<sup>b</sup> | 0.41<sup>c</sup> | ND | 1.40±1.92 |
|  | C18:0 | 16.10<sup>a</sup> | 17.88<sup>b</sup> | 15.03<sup>a</sup> | 15.06<sup>a</sup> | 12.78<sup>c</sup> | 15.37±1.85 |
|  | C20:0 | 3.67<sup>a</sup> | 3.78<sup>a</sup> | 1.98<sup>b</sup> | 2.49<sup>c</sup> | ND | 2.98±0.89 |
| MFA | Total | 43.79 | 31.73 | 39.88 | 37.51 | 26.18 | 35.82±6.94 |
|  | C15:1 | 1.07<sup>a</sup> | 0.73<sup>b</sup> | 1.33<sup>c</sup> | 0.96<sup>a</sup> | ND | 1.02±0.25 |
|  | C16:1 | 36.74<sup>ac</sup> | 30.98<sup>b</sup> | 37.48<sup>c</sup> | 35.64<sup>a</sup> | 26.18<sup>c</sup> | 33.40±4.76 |
|  | C17:1 | 5.98<sup>a</sup> | ND | 1.05<sup>b</sup> | 0.91<sup>b</sup> | ND | 2.65±2.89 |
| PUFA | Total | 30.76 | 24.68 | 24.79 | 32.49 | 41.91 | 30.93±7.07 |
|  | C18:2n6 | 6.78<sup>a</sup> | 13.72<sup>b</sup> | 3.64<sup>c</sup> | 4.99<sup>d</sup> | 9.39<sup>a</sup> | 7.70±3.99 |
|  | C20:3n3 | 4.26<sup>a</sup> | 2.22<sup>b</sup> | 3.63<sup>c</sup> | 3.88<sup>ac</sup> | 5.75<sup>d</sup> | 3.95±1.27 |
|  | C20:5n3 | 3.35<sup>a</sup> | 1.61<sup>b</sup> | 2.24<sup>c</sup> | 4.15<sup>d</sup> | 3.14<sup>a</sup> | 2.90±0.99 |
|  | C22:6n3 | 18.31<sup>a</sup> | 7.13<sup>b</sup> | 15.28<sup>c</sup> | 19.44<sup>a</sup> | 23.62<sup>d</sup> | 16.76±6.16 |

C14:0 (myristic acid), C18:0 (stearic acid), C20:0 (archidic acid), C15:1 (cis-10-pentadecenoic acid), C16:1 (palmitoleic acid), C17:1 (cis-10-heptadecenoic acid), C18:2n6 (linoleic acid), C20:3n3 (cos-11,14,17-eicosatrienoic acid). C20:5n3 (eicosapentaenoic acid), C22:6n3 (docosahexaenoic acid). SFA: saturated fatty acids, MFA: monounsaturated fatty acids, PUFA: polyunsaturated fatty acids. Rows with the same superscript are statistically equals according to the Tuckey's test.

#### 3.1.4 BFT in vitro digestibility

BFT in vitro digestibility did not showed significant differences among the analyzed samples. The calibration curve from the analysis of standard casein had an R^2^ =0.99. Using the line equation from the calibration curve (*y* = 0.001*x* + 0.0117) we determined the protein concentration through all the digestive process phases for samples BFT1 and BFT3. The amount of digested protein in BFT1 was 93.71% (Table 4)

**Table 4.** Concentration of digested BFT protein.

| Phase | ABS (UA) | Concentration (mg/mL) | Average | Std. Dev. | Variance |
| --- | --- | --- | --- | --- | --- |
| t0 | 0.05 | 44.4 | 41.35 | 4.31 | 10% |
| t0 | 0.05 | 38.3 |  |  |  |
| t0E | 0.06 | 48.8 | 42.55 | 8.83 | 21% |
| t0E | 0.04 | 36.3 |  |  |  |
| GP30 | 0.06 | 56.0 | 52.30 | 5.23 | 10% |
| GP30 | 0.06 | 48.6 |  |  |  |
| GP60 | 0.07 | 62.9 | 61.85 | 1.48 | 2% |
| GP60 | 0.07 | 60.8 |  |  |  |
| IP0 | 0.08 | 78.2 | 69.95 | 11.66 | 17% |
| IP0 | 0.07 | 61.7 |  |  |  |
| IP30 | 0.07 | 66.8 | 68.75 | 2.75 | 4% |
| IP30 | 0.08 | 70.7 |  |  |  |
| IP120 | 0.08 | 77.1 | 88.75 | 12.23 | 14%` |
| IP120 | 0.10 | 94.4 |  |  |  |

t0: Initial time, t0E: Zero time with enzyme addition, PG30: gastric phase at 30 minutes, PG60: gastric phase at 60 minutes, IP0: Intestinal phase at time zero with enzymes, IP30: Intestinal phase at 30 minutes, IP120: Intestinal phase at 120 minutes.

### 3.2 BFT population analyses

#### 3.2.1 Data mining

DNA sequences used in this study were mined from BOLD system (Barcode of Life Data System) and from the NCBI (National Center Biotechnology Information). We obtained 106 COI sequences (Supplementary table 1) and five sequences of this study, 111 sequences in total. All COI sequences were trimmed to 622 bp and grouped by geographical region as Atlantic-North(AtlN) (n= 2), Gulf of Mexico (GdM) (n=11), Caribbean Sea (MC) (n= 91), Atlantic-Center (AtlC) (n=1) and Atlantic-South (AtlS) (n=6) (Figure 1). The CR analysis was composed of 14 BFT sequences of 283 bp, five from the Gulf of Mexico and nine from NCBI.

**Figure 1.**
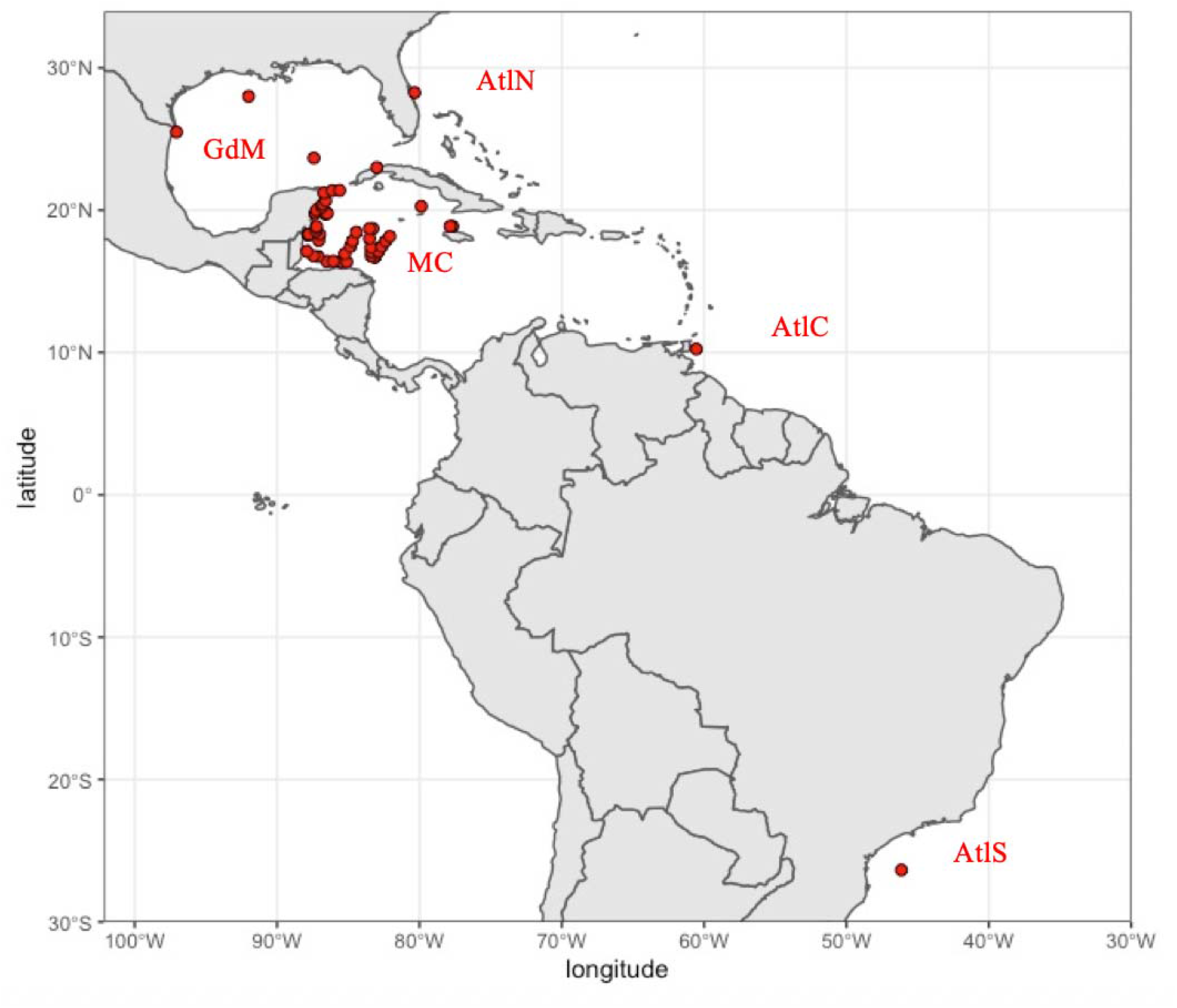
Geographical origins of analyzed samples. GdM: Gulf of Mexico, AtlN: North Atlantic Ocean, MC: Caribbean Sea, AtlC: Central Atlantic Ocean, AtlS: South Atlantic Ocean.

#### 3.2.2. Genetic diversity and haplotype network

BFT COI from the Gulf of Mexico was sequenced and deposited in NCBI under accession numbers MT871943, MT871944, MT871945, MT871946, MT871947 and MT871948. The genetic diversity analysis showed a nucleotide diversity of 0.001, 24 segregating sites and 10 parsimony sites. We observed that the nucleotide diversity was lower to the expected value and in consequence, Tajima’s D is negative (−2.30; P=0.004). After calculate the value of Tajima’s D by region only Mc was significant (−2.32; P=0.01). The median joining network exhibited two main haplotypes (Figure 2), the H5, found in four geographical zones (AtlN, GdM, and Mc) and the H3 found within three geographical zones (GdM, Mc, AtlC), 22 total haplotypes, 6 shared between two or more geographical zones and 14 unique haplotypes (11 found in the Caribbean Sea, 1 in the Gulf of Mexico and 2 in the Atlantic-South).

**Figure 2.**
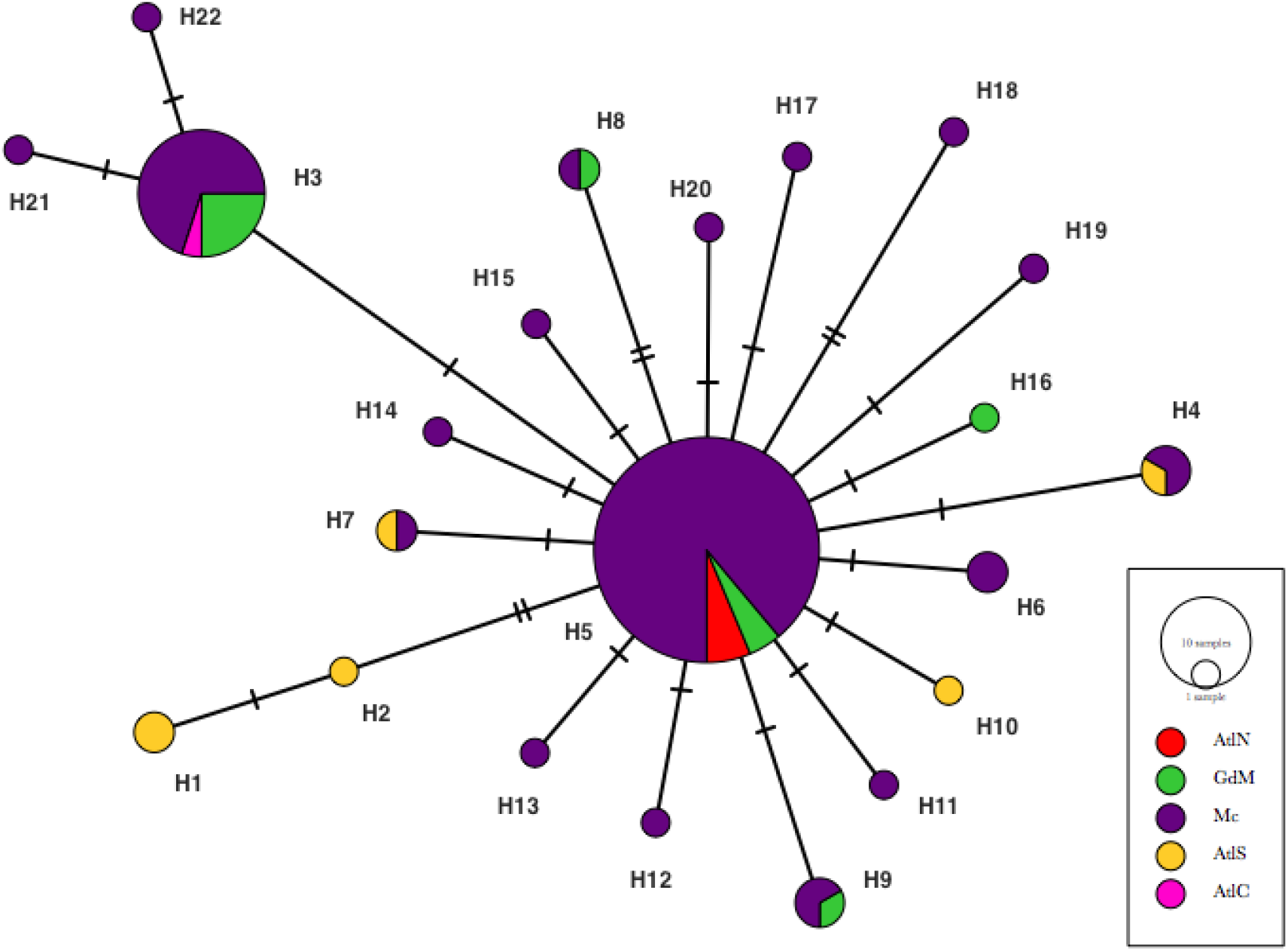
Haplotype network of BFT COI. The color represents the geographical locations of the samples. The circle size is proportional to the haplotype frequency and the dashes on branches represents the number of mutations between nodes (AtlN: North Atlantic, GdM: Gulf of Mexico, Mc: Caribbean Sea, AtlS: South Atlantic, AtlC: Central Atlantic). The smallest circles are unique haplotypes.

**Figure 3.**
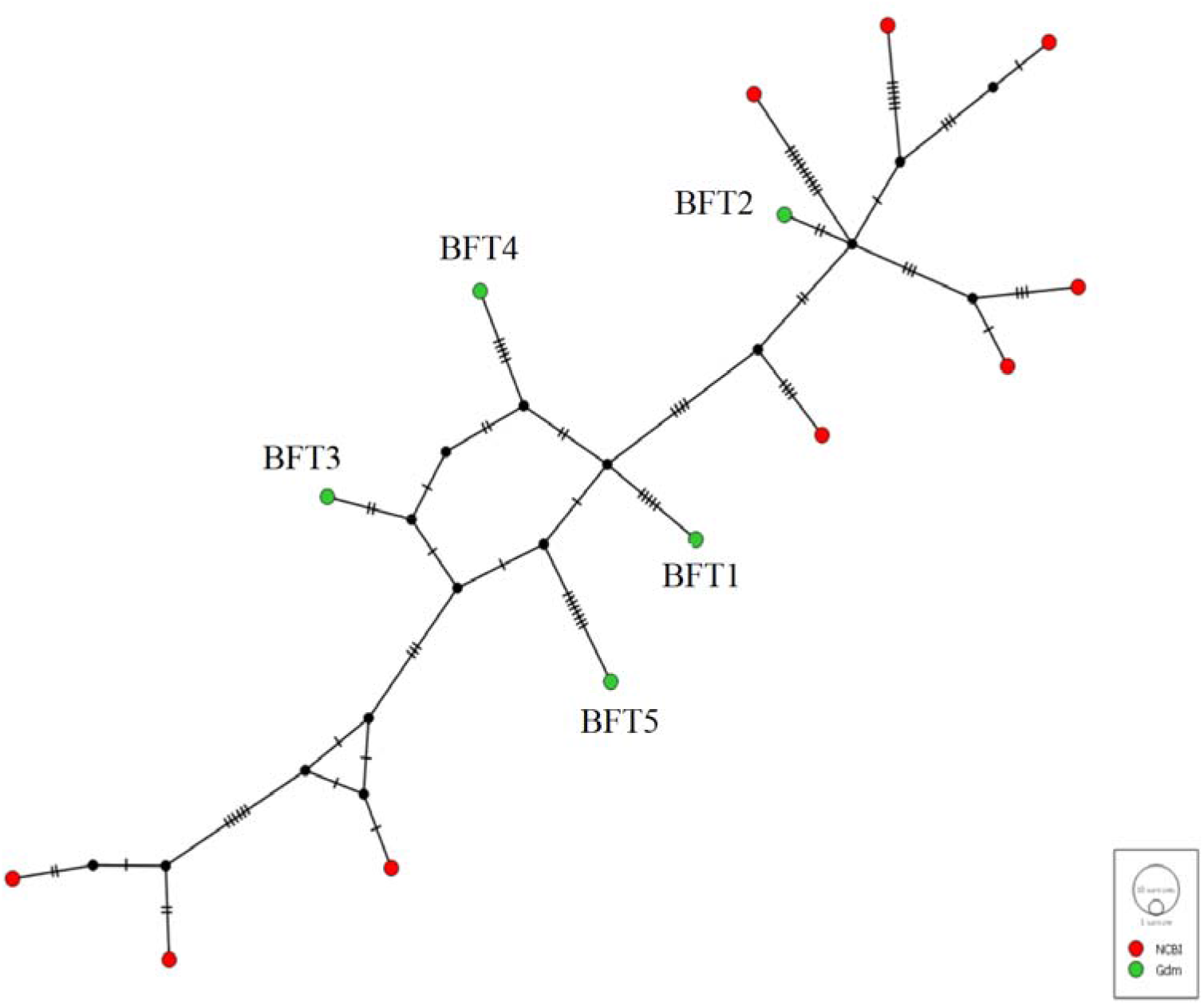
Haplotype network for BFT CR. The color represents the origin of the samples, BFT from GdM are represented in green while NCBI samples are represented in red. The circle size is proportional to the haplotype frequency and the dashes on branches represents the number of mutations between nodes.

The sequences of the CR of the five GdM BFT samples were submitted to the BOLD System Database under IDs THUNN001-20, THUNN002-20, THUNN003-20, THUNN004-20 and THUNN005-20. For the CR, we found a nucleotide diversity of 0.049, 53 segregating sites, 23 parsimony sites. Only 14 unique haplotypes were found for the CR. Although Tajima’s D value was negative (−0.7826) the neutral hypothesis cannot be rejected (P=0.228).

#### 3.2.3 Statistical analysis (AMOVA)

The genetic structure analysis (Table 5) found that the highest variance of BFT COI was 78.92% within each region while the variance among the five geographical locations analyzed was 21.08%. The genetic structure test between the Atlantic Ocean North and Central geographical regions, the Gulf of Mexico and the Caribbean Sea with respect to Atlantic South (AtlN, AtlC, GdM, MC / AtlS) showed the highest variance within each geographical zone (57.84%) but remarkably, the variance among regions increased (42.16%). Finally, the last structure made from two clades (H5 and H3) showed that the highest variance occurred among clades (60.66%).

**Table 5.** Analysis of molecular variance (AMOVA) for COI and CR.

| Sources of variation | DF | Sum of squares | Variance components | % variation |
| --- | --- | --- | --- | --- |
| COI among geographical regions |  |  |  |  |
| Among AtlN, GdM, MC, AtlS, AtlC | 4 | 5.9220 | 0.1186 | 21.08 |
| Within geographical zones | 106 | 47.0870 | 0.4442 | 78.92 |
| Total | 110 | 53.0090 | 0.5628 |  |
| $F_{ST} = 0.2107$ (P=0.0006) | | | | |
| COI two groups |  |  |  |  |
| AtlN, GdM, MC, AtlC/AtlS | 1 | 4.157 | 0.3267 | 42.16 |
| Within populations | 109 | 48.852 | 0.4481 | 57.84 |
| Total | 110 | 53.009 | 0.7748 |  |
| $F_{ST} = 0.4216$ (P=0.0000) | | | | |
| COI two clades |  |  |  |  |
| Clade H5, clade H3 | 1 | 17.8640 | 0.4972 | 60.66 |
| Within populations | 109 | 35.1450 | 0.3224 | 39.34 |
| Total | 110 | 53.0090 | 0.8196 |  |
| $F_{ST} = 0.6066$ (P=0.0000) | | | | |

Geographical regions, AtlN: North Atlantic, GdM: Gulf of Mexico, MC: Caribbean Sea, AtlS: South Atlantic, AtlC: Central Atlantic.

#### 3.2.4 Neothunnus phylogenetic analysis

*Neothunnus* phylogeny was reconstructed from COI with 147 sequences of 622 bp and CR with 111 of 283 bp long. For the COI sequences, the topology of the trees groups *T. tonggol* as one single monophyletic clade whereas BFT, *T. obesus* and *T. albacares* do not show a clear monophyly (Figure 4). On the other hand, the CR showed BFT forming a single monophyletic cluster whereas *T. tonggol* had low support values and *T. albacares* do not group as a single cluster (Figure 5).

**Figure 4.**
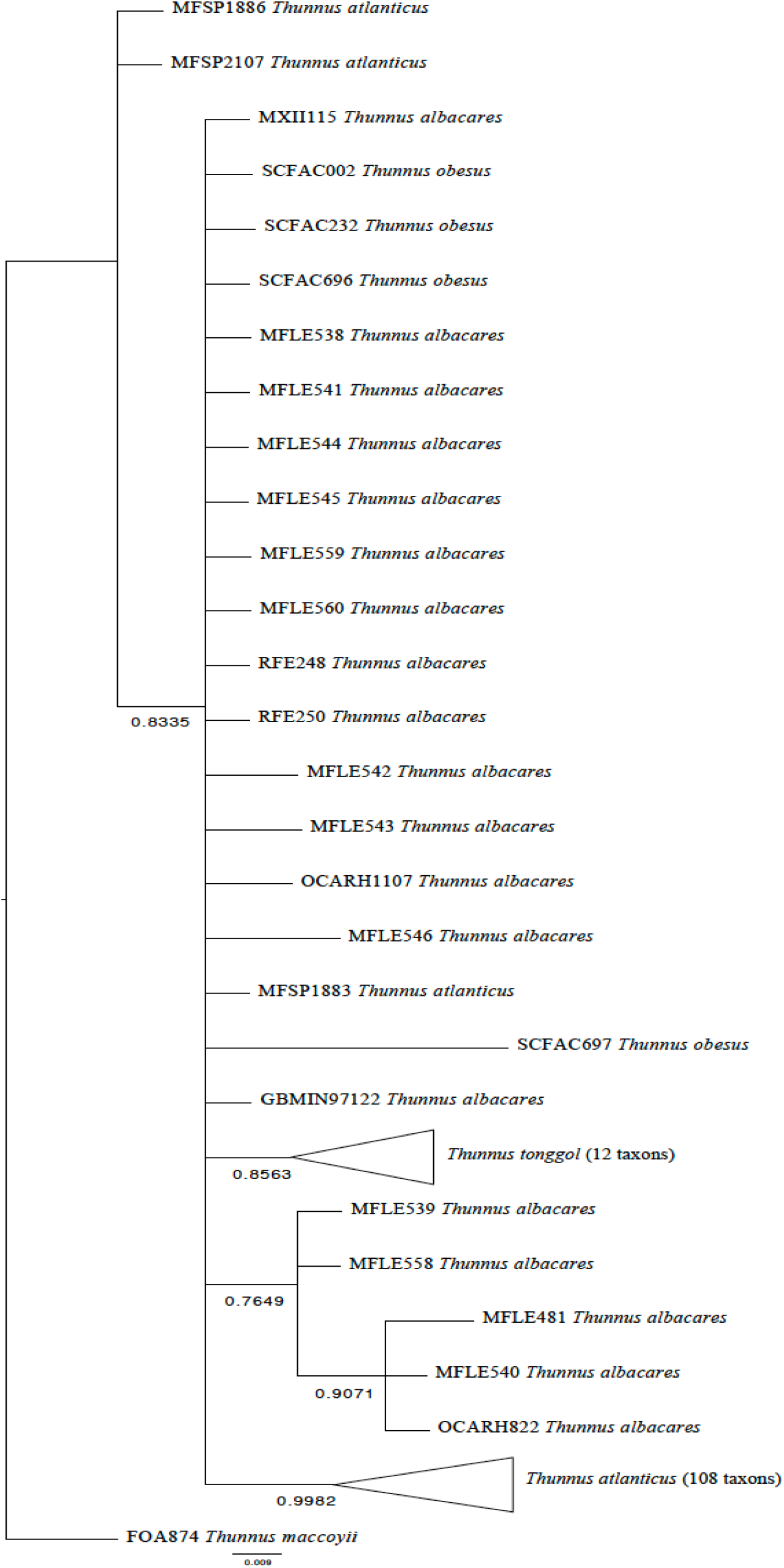
COI Bayesian phylogenetic relationships within the Neothunnus subgenus.

**Figure 5.**
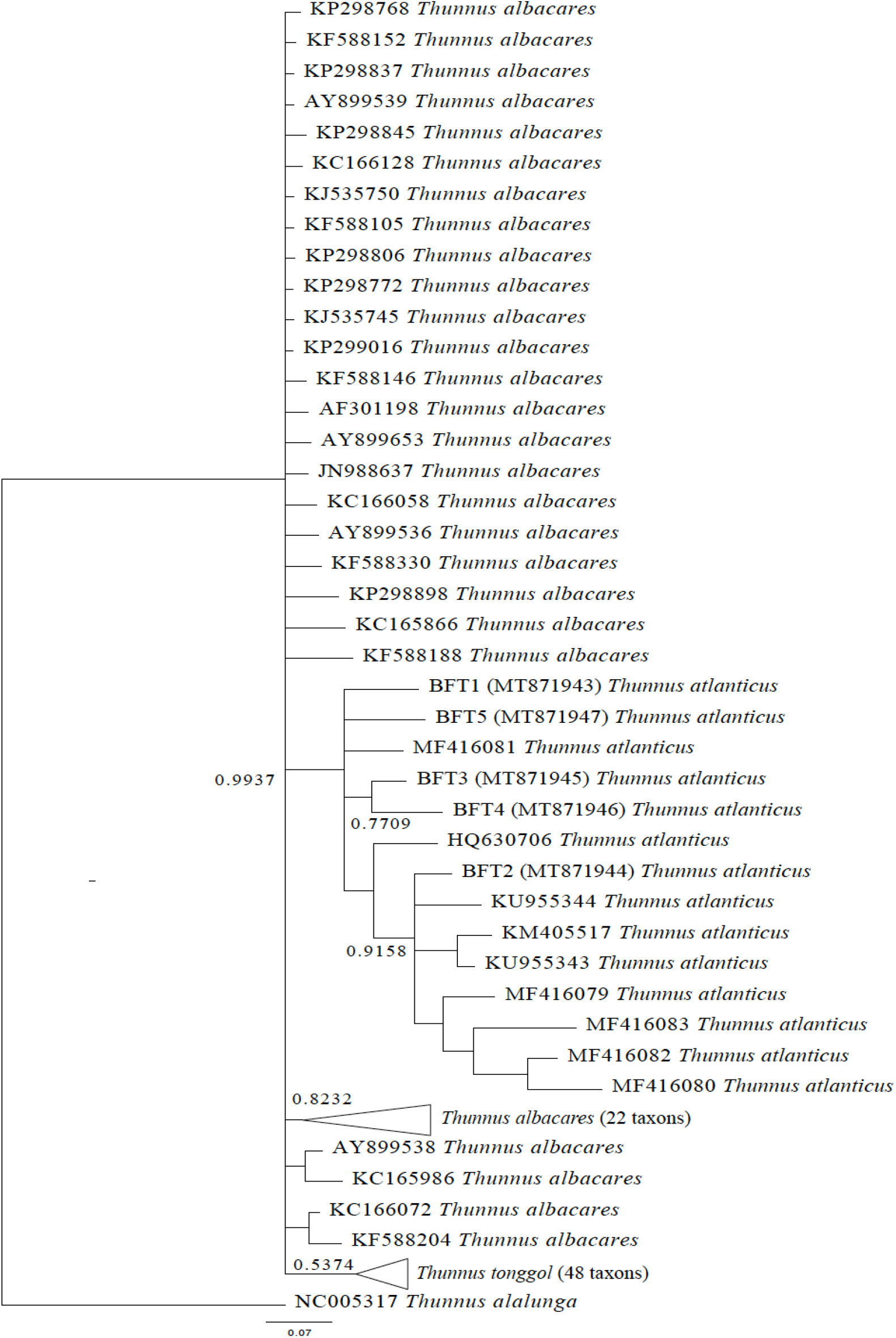
CR Bayesian phylogenetic relationships within the Neothunnus subgenus.

## 4. DISCUSSION

### 4.1 GdM BFT proximate analyses

Fish meat consumption is highly recommended for its nutritional properties and is considered a key part in a balanced healthy diet. Among the quality properties found in fish meat are a low content in fat, high quantities of protein and the presence of trace minerals (Hernandez-Martinez et al., 2015). Fish is one of the most important sources of polyunsaturated fatty acids (PUFA) which are associated with prevention of cardiovascular disease (CVD) due their role in reduction of plasma cholesterol and triglyceride levels (Bird et al., 2018). In this study, we performed the proximate analysis of BFT in order to determine the potential nutritional value of this unexploited species from the American Atlantic coast compared to commercial tuna species. The analyzed BFT caught in the Gulf of Mexico showed an average of 74±5.8 cm of body length and 6.3±0.9 kg of weight, those measurements correspond to the physical and morphological characteristics of BFT that ranges from 30-100 cm and 5 to 7 kg respectively (Griffin et al., 2014, Taquet et al., 2000). In this study, we determined moisture, ash, fatty acids, protein and carbohydrates of BFT. There are extensive studies on nutritional properties in commercial tuna species such as the yellowfin tuna (*T. albacares*) and the bluefin tuna (*T. thynnus*) although in our knowledge this is the first study focused on the nutritional properties of BFT. Compared with commercial species, we found that BFT do not have significant differences in moisture (71-75%) and ash (1.1-1.5%). BFT dry basis protein percentage (86-96%) was significantly higher than reported in *T. thynnus, T. alalunga, orientalis* and *T. albacares* (P<0.001) (Castrillo N et al., 1996, Izquierdo et al., 2007, Karunarathna and Attygalle, 2010, Mohanty et al., 2019, Murase and Saito, 1996, Nurjanah et al., 2020, Vileg and Murray, 1988, Wheeler and Morrisey, 2003).

The dry basis content of total fat BFT found in average was 1.4%, this content is lower than the fat content reported in *T. orientalis* (P=0.0057) and *T. thynnus* (P=0.0094) (Figure 6). However, total fat content is greater in farmed fish (yellowfin tuna and bluefin tuna) than in wild fish (P=0.001), this could be explained by the type of diet given to farmed fish (Casalduero and Jerez, 2006, Estess et al., 2014, Nakamura et al., 2007, Roy et al., 2010, Topic Popovic et al., 2012, Vizzini et al., 2010) Besides the total fat content in BFT, we analyzed those fatty acids with human health implications, dietary omega 3 (n3), and omega 6 (n3) polyunsaturated fatty acids. We observed that BFT showed an average content of eicosapentaeonic acid (EPA) (n3) of 2.90±0.99 and an average of docosahexaenoic acid (DHA) (n3) of 11.17±8.31. A previous report from Cai et al. (2007) in fishes from the Northern Gulf of Mexico demonstrated that the content of EPA and DHA in BFT is 6.35% and 23.63%, in this context, the low content of EPA and DHA in the analyzed BFT samples could be evidence of genetic structure within the Gulf of Mexico.

**Figure 6.**
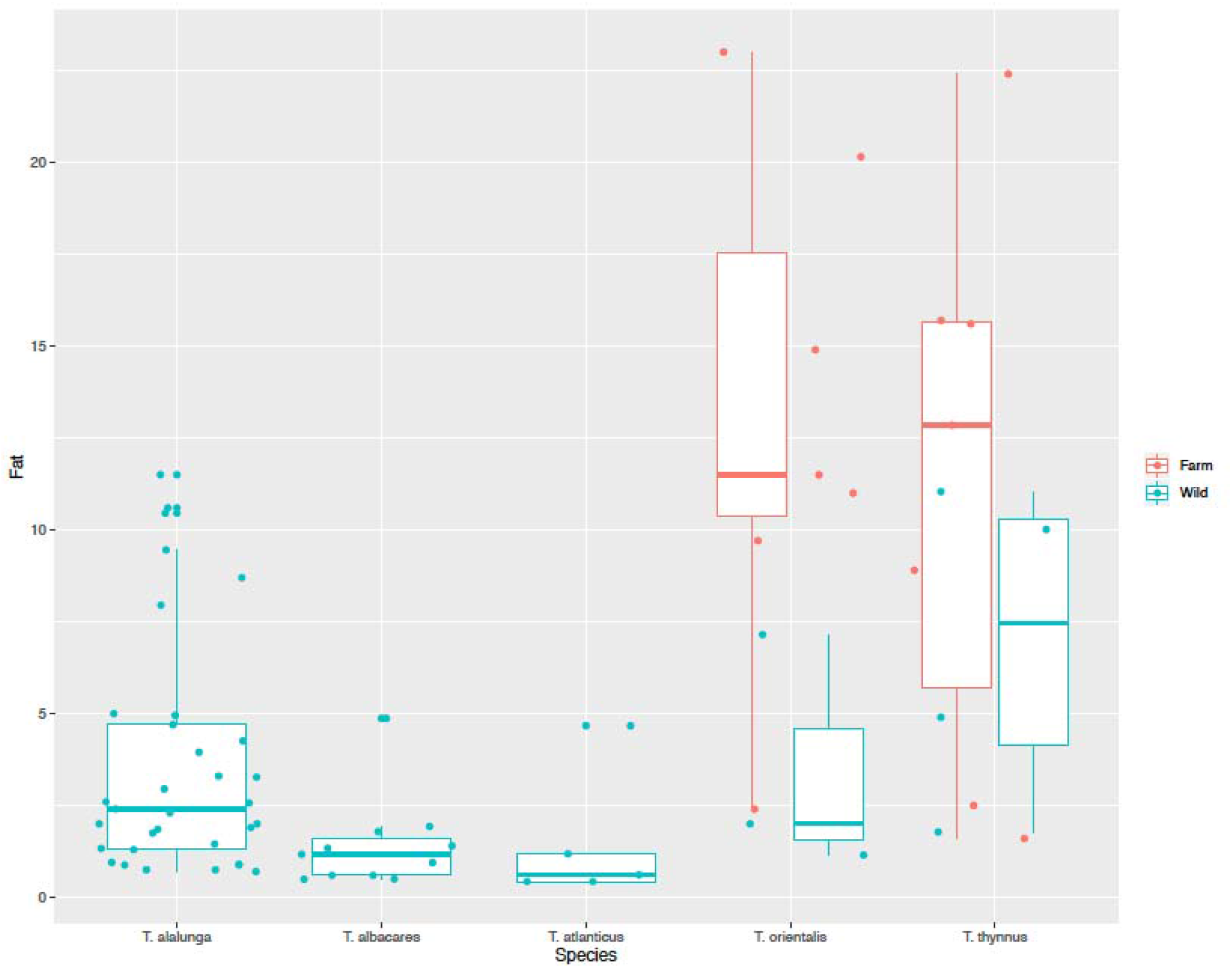
Total fat content in *T. alalunga, T. albacares*, BFT, *T. orientalis* and *T. thynnus*

DHA levels in BFT (16.76±6.16) are higher than values reported in wild salmon (12.5±0.22), but lower than values found in yellowfin tuna (YFT) (36.72 ± 1.11) and Atlantic bluefin tuna (ABT) (24-40 %), whereas (-linoleic acid levels are similar in BFT (3.64–13.72 %), ABT (1.16-1.43 %) and salmon 18:2n-6 (1.7–15.1 %) but lower in EPA (7.07 ± 0.09 in YFT, 4.50-5.37 % in ABT and 6.5 ± 0.8 in salmon and 2.9 ± 0.99 in BFT) (Hernandez-Martinez et al., 2015, Jensen et al., 2012, Sprague et al., 2020).

Content of analyzed SFA in BFT (C14:0, C18:0 and C20:0) was found similar in C14:0 and C20:0 to the lipid profile reported in ABT and YFT, although stearic acid (C18:0) is significantly higher in BFT (15.37 ± 1.85) compared with YFT (3.42 ± 0.51) (Renuka et al., 2016). Evaluation of BFT flesh quality with respect to fatty acid contents in this study demonstrated that this species has a comparable nutritional composition to similar commercial species widely available in Mexico and other countries and represent a viable source of PUFA and protein for human consumption.

### 4.2 BFT population analysis

Sequences analyzed in this research were obtained from the Canadian database BOLD System, it was originally created to store COI gene data which is considered a barcode in animals (Ratnasingham and Hebert, 2007). Previous comparative studies in fish concluded that COI have limitation for species identification, but the hypervariable CR allows correct identification of species within the *Thunnus* genus. We found in COI a nucleotide diversity similar to the reported by Viñas and Tudela (2009) (π=0.001) whereas the CR, the diversity (π=0.049) was higher to the diversity reported for BFT in the Atlantic Ocean and the Gulf of Mexico (Chow et al., 2006, Saxton, 2009). analyzed the genetic variations in a 323 bp region of the COI and six microsatellite loci (msat). The results suggest the presence of two genetically distinct populations in the Western Atlantic Ocean (msat st=0.01, P=0.006 and mtDNAst=0.01, P=0.049) between the GdM and the AtlN. In our study, within the five geographical zones included, we found a total 22 haplotypes of BFT. 12 were exclusively found in the Caribbean Sea, one identified in the Gulf of Mexico and three in Brazil (South Atlantic Ocean). Six haplotypes were shared among the studied zones. This could be relevant for BFT stocks identification since there are reports of spawning sites in the Saint Peter and Saint Paul archipelagos in Brazil (Bezerra et al., 2013). Other spawning and feeding sites are located in the Caribbean Sea and the Banco Chinchorro in the North Atlantic Ocean (Fenton et al., 2015, Luckhurst et al., 2001). The genetic structure of the found haplotypes reveals that the highest variance occurred within the analyzed regions. However, variance in the distribution zones was high comparing North-Central Atlantic, MC and GdM specimens with South Atlantic (42.16%), this is evidence that the genetic flow among these regions is limited., Additional studies with nuclear markers may be helpful describing migration patterns of the BFT from the Caribbean Sea to the Atlantic South since the haplotypes from Brazil were not found beyond this region. For the CR, we could not stablish groups due the low number of sequences reported in the databases. Although BFT is a minor concern for conservationists there is a very limited knowledge of their population size, Tajima D indicated an expansion scenario for COI in the Caribbean Sea, it is probably due to different cohorts and the excess of unique haplotypes. In the CR was not significant for expansion. In this context, several studies in the Gulf of Mexico demonstrated that at larval stage BFT is the pelagic species most common of Scombridae (Cornic and Rooker, 2018, Pruzinsky et al., 2020).

Due the limited information provided by the COI to identify species within the *Thunnus* genus, the CR have been recently used, since it is more informative even than the ITS1 (Viñas and Tudela, 2009). Another factor influencing the correct phylogenetic reconstruction in the *Thunnus* genus is the species hybridization. In BFT there are no evidence nor reports of hybridization events with any species in the subgenus *Neothunnus* despite the close phylogenetic relationships between the species. Previous studies located *T. obesus* within the *Thunnus* subgenus because its adaptation to template waters, nevertheless, whole genome sequencing and transcriptomic data concur that *T. obesus* is closely related to the species within the *Neothunnus* subgenus (Ciezarek et al., 2019, Díaz-Arce et al., 2016). In our phylogenetic analysis, we included *T. obesus* in order to reconstruct the Neothunnus phylogeny. Our COI results concluded that only *T. tonggol* can be identified as a single cluster species and BFT have a main cluster but two Brazilian sequences (MFSP1886, MFSP2107) were clustered separately. One sequence from the South Atlantic (MFSP1883) exhibited a paraphyletic cluster together with *T. obesus* and *T. albacares*. CR phylogeny also identified BFT as a monophyletic group. However, *T. tonggol, T. obesus* and *T. albacares* clades had low statistical support.

## ACKNOWLEDGMENTS

We thank the Mexican Consejo Nacional de Ciencia y Tecnología for the support given to this project, the IPN-SIP research grants given to Nadia A. Fernández Santos (20201972) and Xochitl F. De la Rosa (20196142). Also we thank Dr. Mario A. Rodríguez, Dr. Ana Luisa González-Pérez and Dr. Guadalupe C. Rodríguez-Castillejos for their valuable recommendations and review.

## DECLARATION OF COMETING INTEREST

The authors report no competing declarations of interest.

## AUTHOR CONTRIBUTIONS

**Conceptualization:** XFDRR, ALPT, HMM.

**Funding acquisition:** XFDRR, ALPT, JAVR, NAFS, HMM.

**Investigation:** JRRR, XFDRR, NAFS, YMNM, ALPT, JAVR, HMM.

**Writing – original draft:** XFDRR, HMM.

**Writing – manuscript review and edition:** XFDRR, ALPT, JAVR, NAFS, HMM

