## Supplemental Table 1 for "Proximate and Genetic Analysis of Blackfin Tuna (*T. atlanticus*)": BFT-Table-S1.docx

Table S1. Accession numbers of COI sources for phylogenetic analyses

| Sample ID | Locality | Latitude/Longitude | Distribution Zone |
| --- | --- | --- | --- |
| BIOUG30362_G1 | Mexico | 20.66,-86.5898 | Caribbean Sea |
| BIOUG30362_H9 | Mexico | 20.66,-86.5898 | Caribbean Sea |
| BIOUG30479_E02 | Mexico | 19.714,-86.5992 | Caribbean Sea |
| BIOUG30479_G07 | Mexico | 19.714,-86.5992 | Caribbean Sea |
| BIOUG30479_G10 | Mexico | 19.714,-86.5992 | Caribbean Sea |
| CH-0000664_F10 | Cuba | 23,-83 | Atlantic |
| CH-0000664_F12 | Cuba | 23,-83 | Atlantic |
| BW-A1162 | United states |  | Atlantic-North |
| BW-A1164 | United states |  | Atlantic-North |
| BW-A1165 | United states |  | Gulf of Mexico |
| BW-A1166 | United states |  | Gulf of Mexico |
| MFL281 | Mexico | 18.283,-87.819 | Caribbean Sea |
| MFL282 | Mexico | 18.283,-87.819 | Caribbean Sea |
| MFL284 | Mexico | 18.283,-87.819 | Caribbean Sea |
| MFL319 | Mexico | 18.28,-87.82 | Caribbean Sea |
| MFL320 | Mexico | 18.28,-87.82 | Caribbean Sea |
| MFL246 | Mexico | 18.28,-87.82 | Caribbean Sea |
| MFL247 | Mexico | 18.28,-87.82 | Caribbean Sea |
| MFL248 | Mexico | 18.28,-87.82 | Caribbean Sea |
| MFL249 | Mexico | 18.28,-87.82 | Caribbean Sea |
| MFL250 | Mexico | 18.28,-87.82 | Caribbean Sea |
| MFLE4651 | Jamaica | 18.869,-77.665 | Caribbean Sea |
| MFLE4740 | Jamaica | 18.873,-77.845 | Caribbean Sea |
| MFLE4741 | Jamaica | 18.873,-77.845 | Caribbean Sea |
| MFLE4819 | Honduras | 16.945,-85.248 | Caribbean Sea |
| MFLE4936 | Mexico | 21.373,-86.138 | Caribbean Sea |
| MFLE4937 | Mexico | 21.388,-85.598 | Caribbean Sea |
| MFLE4954 | Mexico | 20.233,-86.85 | Caribbean Sea |
| MFLE4962 | Mexico | 19.783,-86.47 | Caribbean Sea |
| MFLE4969 | Mexico | 20,-87.25 | Caribbean Sea |
| MFLE4971 | Mexico | 19.767,-87.336 | Caribbean Sea |
| MFLE4973 | Mexico | 19.767,-87336 | Caribbean Sea |
| MFLE4975 | Mexico | 19.767,-87336 | Caribbean Sea |
| MFLE4978 | Mexico | 19.349,-87.001 | Caribbean Sea |
| MFLE4981 | Mexico | 18.86,-87.544 | Caribbean Sea |
| MFLE4983 | Mexico | 18.862,-87.228 | Caribbean Sea |
| MFLE5001 | Mexico | 18.784,-87.257 | Caribbean Sea |
| MFLE5014 | Mexico | 18.309,-87.771 | Caribbean Sea |
| MFLE5015 | Mexico | 18.312,-87.003 | Caribbean Sea |
| MFLE5021 | Mexico | 18.531,-87268 | Caribbean Sea |
| MFLE5024 | Mexico | 23.667,-87.417 | Gulf of Mexico |
| MFLE5027 | Mexico | 23.667,-87.417 | Gulf of Mexico |
| MFLE5028 | Mexico | 23.667,-87.417 | Gulf of Mexico |
| MFLE5029 | Mexico | 23.667,-87.417 | Gulf of Mexico |
| MFLE5032 | Atlantic Ocean | 18.706,-83.263 | Caribbean Sea |
| MFLE5033 | Atlantic Ocean | 18.706,-83.263 | Caribbean Sea |
| MFLE5034 | Atlantic Ocean | 18.161,-82.078 | Caribbean Sea |
| MFLE5035 | Honduras | 17.1,-82.872 | Caribbean Sea |
| MFLE5037 | Honduras | 16.701,-83.132 | Caribbean Sea |
| MFLE5039 | Honduras | 16.365,-85532 | Caribbean Sea |
| MFLE5041 | Honduras | 16.409,-86.512 | Caribbean Sea |
| MFLE5043 | Belize | 16.409,-86.512 | Caribbean Sea |
| MFLE5044 | Belize | 17.399,-86.677 | Caribbean Sea |
| MFLE5045 | Belize | 17.399,-86.677 | Caribbean Sea |
| MFLE5046 | Belize | 17.399,-86.677 | Caribbean Sea |
| MFLE5047 | Belize | 16.744,-87.118 | Caribbean Sea |
| MFLE5048 | Belize | 16.744,-87.118 | Caribbean Sea |
| MFLE5051 | Belize | 17.876,-87.067 | Caribbean Sea |
| MFLE5052 | Belize | 16.783,-87.445 | Caribbean Sea |
| MFLE5053 | Cayman Islands | 17.108,-87.913 | Caribbean Sea |
| MFLE5054 | Cayman Islands | 20.277,-79.867 | Caribbean Sea |
| MFLE5056 | Honduras | 18.706,-83.263 | Caribbean Sea |
| MFLE5057 | Atlantic Ocean | 17.45,-82.649 | Caribbean Sea |
| MFLE5058 | Honduras | 17.832,-82.38 | Caribbean Sea |
| MFLE5059 | Honduras | 17.45,-82.649 | Caribbean Sea |
| MFLE5060 | Honduras | 16.701,-83.132 | Caribbean Sea |
| MFLE5061 | Atlantic Ocean | 16.701,-83.132 | Caribbean Sea |
| MFLE5062 | Atlantic Ocean | 18.706,-83.263 | Caribbean Sea |
| MFLE5063 | Atlantic Ocean | 18.706,-83.263 | Caribbean Sea |
| MFLE5064 | Honduras | 18.706,-83.263 | Caribbean Sea |
| MFLE5065 | Honduras | 16.773,-83.391 | Caribbean Sea |
| MFLE5066 | Honduras | 16.773,-83.391 | Caribbean Sea |
| MFLE5067 | Honduras | 16.773,-83.391 | Caribbean Sea |
| MFLE5068 | Honduras | 17.064,-83.389 | Caribbean Sea |
| MFLE5069 | Honduras | 17.064,-83.389 | Caribbean Sea |
| MFLE5070 | Honduras | 17.396,-83.401 | Caribbean Sea |
| MFLE5071 | Honduras | 17.396,-83.401 | Caribbean Sea |
| MFLE5072 | Honduras | 17.396,-83.401 | Caribbean Sea |
| MFLE5073 | Honduras | 18.015,-83.51 | Caribbean Sea |
| MFLE5074 | Honduras | 18.015,-83.51 | Caribbean Sea |
| MFLE5075 | Honduras | 18.015,-83.51 | Caribbean Sea |
| MFLE5076 | Honduras | 18.015,-83.51 | Caribbean Sea |
| MFLE5077 | Atlantic Ocean | 18.015,-83.51 | Caribbean Sea |
| MFLE5078 | Atlantic Ocean | 18.708,-83.512 | Caribbean Sea |
| MFLE5079 | Honduras | 18.452,-84.44 | Caribbean Sea |
| MFLE5080 | Honduras | 17.838,-84.699 | Caribbean Sea |
| MFLE5081 | Honduras | 17.838,-84.699 | Caribbean Sea |
| MFLE5082 | Honduras | 17.45,-84.844 | Caribbean Sea |
| MFLE5083 | Honduras | 16.374,-85.103 | Caribbean Sea |
| MFLE5084 | Honduras | 16.42,-86.04 | Caribbean Sea |
| MFLE5085 | Honduras | 16.42,-86.04 | Caribbean Sea |
| LBPV53027 | Brazil | -26.352,46.139 | Atlantic-South |
| LBPV53028 | Brazil | -26.352,-46.139 | Atlantic-South |
| LBPV53029 | Brazil | -26.352,-46.139 | Atlantic-South |
| LBPV53030 | Brazil | -26.352,-46.139 | Atlantic-South |
| LBPV53031 | Brazil | -26.352,-46.139 | Atlantic-South |
| LBPV53030.1 | Brazil | -26.352,-46.139 | Atlantic-South |
| MX927 | Mexico | 20.416,-86.804 | Caribbean Sea |
| MX931 | Mexico | 20.416,-86.804 | Caribbean Sea |
| MXIII424 | Mexico | 21.22,-86.721 | Caribbean Sea |
| MXIII428 | Mexico | 21.22,-86.721 | Caribbean Sea |
| MXIII432 | Mexico | 21.22,-86.721 | Caribbean Sea |
| MXIII435 | Mexico | 21.22,-86.721 | Caribbean Sea |
| MXIII459 | Mexico | 21.22,-86.721 | Caribbean Sea |
| MXIII460 | Mexico | 21.22,-86.721 | Caribbean Sea |
| TOBA9079 | Trinidad and Tobago | | Atlantic-Center |
